# Algorithmic improvements for discovery of germline copy number variants in next-generation sequencing data

**DOI:** 10.1101/441378

**Authors:** Brendan O’Fallon, Jacob Durtschi, Tracey Lewis, Devin Close

## Abstract

Copy number variants (CNVs) play a significant role in human heredity and disease, however sensitive and specific characterization of CNVs from NGS data has remained challenging. Detection is especially problematic for hybridization-capture data in which read counts are the sole source of copy number information. We describe two algorithmic adaptations that improve CNV detection accuracy in a Hidden Markov Model (HMM) context. First, we present a method for computing target- and copy number state-specific emission distributions. Second, we demonstrate that the Pointwise Maximum *a posteriori* (PMAP) HMM decoding procedure yields improved sensitivity for small CNV calls compared to the more common Viterbi HMM decoder. We develop a prototype implementation, called Cobalt, and compare it to other CNV detection tools using sets of simulated and previously detected CNVs with sizes spanning a single exon up to a full chromosome. In both the simulation and previously detected CNV studies Cobalt shows similar sensitivity but significantly improved positive predictive value (PPV) compared to other callers. Overall sensitivity is 80%-90% for deletion CNVs spanning 1-4 targets and 90%-100% for larger deletion events, while sensitivity is somewhat lower for small duplication CNVs. Cobalt demonstrates significantly improved positive predictive value (PPV) compared to other callers with similar sensitivity, typically making 5X fewer total calls overall.

---

Accurate detection of germline copy number variants (CNVs) from hybridization-capture next generation sequencing (NGS) data remains a significant challenge. The sparse nature of targeted capture data typically precludes detecting CNVs via paired end mapping or split-read analysis, leaving read depth as the primary signal of DNA copy number. Read depth is subject to many confounding factors, including those that affect each region independently, such as GC or CpG content, as well as factors that create correlations in depths between regions, such as the presence of common CNVs or ‘batch effects’ produced by variable lab conditions. Elucidation of the true number of copies of an allele requires accounting for both sources of variability in additional to the stochastic nature of the fragment hybridization process.

Many methods have been proposed to estimate copy-number status from NGS read depth data. Love et al. (2011) directly model read counts at each target with negative binomial distributions, and construct a Hidden Markov Model (HMM) whose parameters are estimated by a maximum likelihood procedure. Li et al. (2012) suggest a method where the ratio of sample to control set read depths is examined. For each target, a mean and variance parameter is estimated from the control set, and sample variation is assumed to follow a normal distribution. More recently, Johannsen et al. (2016) describe a method that examines the ratio of standardized read depths in a sample compared to a subset of samples from control set, selected by a measure of similarity to the test sample. Depth standardization is performed both by taking sample-level Z-scores as well as normalizing depths within genes. CNV calls are produced by collecting targets that exceed a given z-score threshold, with an ad-hoc algorithm that combines both scores.

Of particular relevance here, Krumm et al. (2012) and Fromer et al. (2012) introduced similar techniques that involve decomposing the matrix of depths across control samples using singular value decomposition (SVD). The largest *k* right singular vectors were retained, and the original data projected into the new, lower dimensional space. Finally, this low dimensional reconstruction was subtracted from the original depth matrix, thereby removing major axes of variation and exposing less common variation, such as that produced by CNVs. The effect of GC and other depth-influencing covariates is not explicitly removed by the SVD, but their influence on the mean depth of a target is removed prior to SVD by centering. The method of Krumm et al. 2012 detects CNVs via an ad-hoc segmentation algorithm, while Fromer et al. (2012) construct an heterogeneous HMM with fixed emission distributions and detect CNVs with Viterbi.

Despite ongoing interest in the topic, several recent reviews and comparison papers have documented relatively low sensitivity, reproducibility, and positive predictive value (PPV) in read-depth based CNV detection methods, often in contrast to values claimed in the method papers. For instance, Hong et al. (2016) found that PPV ranged from 7%-60% for the Krumm et al. (2012) method, and 20% - 37% for Fromer et al. (2012), and sensitivity ranged from 5% to 40%, depending on the data set used. Similarly low sensitivities were found by Yao et al. (2017). Tan et al. (2014) also noted high Mendelian error rates in a set of exome trios and poor concordance of CNV calls between callers. Contrasting perspectives can be found in de Ligt (2013) and Yamamoto et al (2016), who found relatively high sensitivity and specificity in larger, multi-gene CNVs using the Krumm (2012) and Fromer (2012) methods.

A potential cause of disagreement between reviews is the size of the CNVs assessed. Large CNVs are likely to span many probe targets, and hence will be easier to detect by most algorithms. Reviews that examine a small set of large CNVs are likely to find relatively high sensitivity and high concordance with array results, while assessments that examine all calls regardless of size seem likely to find lower PPV.

Here, we introduce a method that combines ideas from both Love et al. (2011) and Fromer et al. (2012). Similar to the Fromer (2012) work we use SVD to identify and remove common sources of variation from raw read depth data. We also construct an HMM with target- and state-specific emission distribution parameters in a manner superficially similar to Love et al. (2011), although our method of parameter estimation differs significantly. Finally, in contrast to both the Fromer and Love methods we decode the HMM using the pointwise maximum a posteriori (PMAP) algorithm (Lember & Koloydenko 2013), instead of the more typical Viterbi algorithm, in an effort to maximize the number of correctly called targets. We compare Cobalt to six other detection tools using both simulated and orthogonally detected CNVs of all sizes. We also demonstrate performance improvements and parallelization opportunities resulting from partitioning of the targets into independent groups.

## 1 Methods

We investigate the performance of our method on two different datasets, exomes and a large custom panel. The exome data was used for a simulation analysis, while the custom panel was used to explore accuracy in detecting CNVs previously discovered by array comparative genomic hybridization (aCGH) or multiplex ligation-dependent probe amplification (MLPA). Throughout we compare our method to previous detection tools, including Conifer (Krumm et al. 2012), XHMM (Fromer et al. 2012), Convading (Johannson et al. 2016), ExomeDepth (Plagnol et al. 2012), Clamms (Packer et al. 2015), and Codex (Jiang et al. 2015).

### 1.1 NGS data

70 exomes derived from whole blood were captured using the xGen Exome Research Panel v1.0 probe set from IDT. Exomes were sequenced on 4 separate runs of an Illumina HiSeq 4000 instrument to a mean read depth of approximately 150 reads. Reads were aligned to human reference genome GRCh37 with phiX and decoy sequences included using BWA MEM (v0.7.12, Li 2013). Potential PCR duplicates were identified and marked using Sambamba (Tarasov et al. 2015).

Read counting was performed using the methods recommended by each caller examined, often using either the DepthOfCoverage tool from GATK 3.7 (McKenna et al 2010) or the multicov facility in BEDTools (Quinlan et al. 2010), although ConVaDINg (Johannson et al. 2016) implements read counting internally. For Cobalt, read counting was performed using the PySAM interface to samtools (Li 2009), and the total number of reads overlapping each CNV target (probe) was recorded.

In addition to the exome data, we analyzed 218 samples captured with a custom probe design manufactured by IDT. This large panel targets 4,921 genes with 71,163 probes and has a footprint of 16Mb. These samples were sequenced on 5 runs of a HiSeq 4000 instrument to a mean read depth of approximately 300. Data analysis was identical to that for the exomes.

### 1.2 Simulated CNV generation

Simulated CNVs were introduced into exome BAM files by either removing or duplicating existing reads. CNVs spanning 1, 3 or 10 exons were generated in separate BAM files, see Table 1. CNV locations were chosen uniformly from RefSeq alignments on the autosomes, and read counts were reduced by 50% to create heterozygous deletions and increased by 50% to create heterozygous duplications.

**Table 1:** Number of simulated CNVs by size

| Number of exons | Number of CNVs |
| --- | --- |
| 1 | 1,000 |
| 3 | 300 |
| 10 | 100 |

### 1.3 Cobalt

Our algorithm operates in two phases, a ‘training’ phase where data from control samples is used to create a reusable model, and a ‘prediction’ phase where the model is used in combination with data from a test sample to detect CNVs.

#### 1.3.1 Training

During training, read count depths from multiple samples are used to generate a collection of parameters referred to as a ‘model’. The samples used for training are termed the ‘background’ or ‘control’ samples. Read count data must be provided in a BED-formatted file, with rows corresponding to targets and columns containing read counts for the samples. Targets do not need to be of similar sizes, but should not be overlapping. Our implementation does not assume any particular method of obtaining read counts - both mean target read depth or the total number of reads overlapping a target region are suitable. Under typical use each target corresponds to a single hybridization probe location, although special considerations should be taken if there are multiple overlapping probe locations.

Read depth data are converted into a *n* × *p* matrix *D*, with *n* targets and *p* samples. We then compute *P* such that

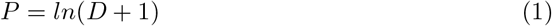

After centering each column of *P* about the column median to produce *R*, we then compute the *k* right singular vectors of *R* via the randomized SVD algorithm of Halko et al. (2011). *k* is computed as the minimum value such that the proportion of variance explained by singular vectors 0..*k* is at least *v*, which by default is 0.90 but can be changed by the user. Setting *C* to be the column matrix of these vectors, we then compute

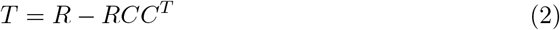

The centering procedure removes target-specific biases due to, for instance, genomic GC or CpG content, while the SVD procedure removes between-target correlations due to CNVs present in the samples or ‘batch effects’ that induce similar changes across sets of targets.

For brevity, we refer to the above steps as *f*, such that *T* = *f* (*P, C*), where *C* is the column matrix of the top *k* right singular vectors of *R*.

#### 1.3.2 Target partitioning

The training procedure above involves computing the matrix *CC^T^*, where *C* is *n* × *k*, which results in an *n × n* matrix. For large numbers of targets (*n*) the amount of storage required may exceed the amount of memory available, and performance may be impacted. To avoid the computation and use of large matrices, we partition the *n* targets into disjoint sets of approximate size *j*, and treat each partition independently. *j* is a user-settable parameter, for the results shown here a value of 1,000 is used. Ad-hoc experiments have suggested that results are generally insensitive to the choice of *j* as long as *j* >> 100. The partitioning algorithm seeks to distribute partitions as uniformly as possible across targets. (Early experiments assigned many adjacent targets to a single partition and had poor sensitivity to CNVs that occupied a large percentage of the partition.)

Target partitioning greatly speeds the training procedure and reduces memory requirements from *O*(*n*^2^) to *O*(*j*^2^) without significantly impacting CNV calling accuracy.

#### 1.3.3 Emission distribution parameter estimation

We assume all emission distributions are Gaussian with means and variances that differ across both states and targets, and we estimate the mean and variance for each state and target separately. Copy number states are described by the expected change in the raw read depth of the target - for instance, for a heterozygous deletion, one might expected a 50% reduction in read depth, while a homozygous duplication may produce a 100% increase in depth. Let vector *s* hold the coefficients describing the expected change in depth associated with each state. Under typical usage, we define 5 states and set *s* = {0.01, 0.5, 1.0, 1.5, 2.0}, where the elements correspond to homozygous deletion, heterozygous deletion, diploid, heterozygous duplication, and homozygous duplication / amplification. Special considerations are made for calls on the X and Y chromosomes in males, as described below.

To estimate the mean and variance parameters for target *t* and state *i*, we *D̂* _*t,i*_ form by multiplying column *t* of *D* by *s_i_*. We then compute 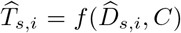, and take the sample mean and sample variance of column *t* of 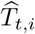 to be the mean and variance of the emission distribution for target *t* and state *i*. The above procedure is repeated for every target and copy number state and the resulting means and variances recorded for all states and targets. We refer to the full collection of parameter estimates and singular vectors *C* as a ‘model’, which is persisted and used to facilitate the CNV discovery procedure.

#### 1.3.4 Target resolution calculation

Using the set of stored emission distribution parameters it is possible to calculate an ad-hoc value that reflects the power of a given target to resolve copy number status in general. For some targets emission distributions are well separated and have small variances relative to the difference in means between states (e.g. figure 1a), while at other targets the distributions may overlap substantially (figure 1c). CNV identification is likely to be impaired when the distributions overlap because posteriors for the states are also likely to overlap substantially. In some cases, it may be possible to detect certain CNV states, such as homozygous deletions, while resolution for other states may be relatively poor (figure 1c).

**Figure 1:**
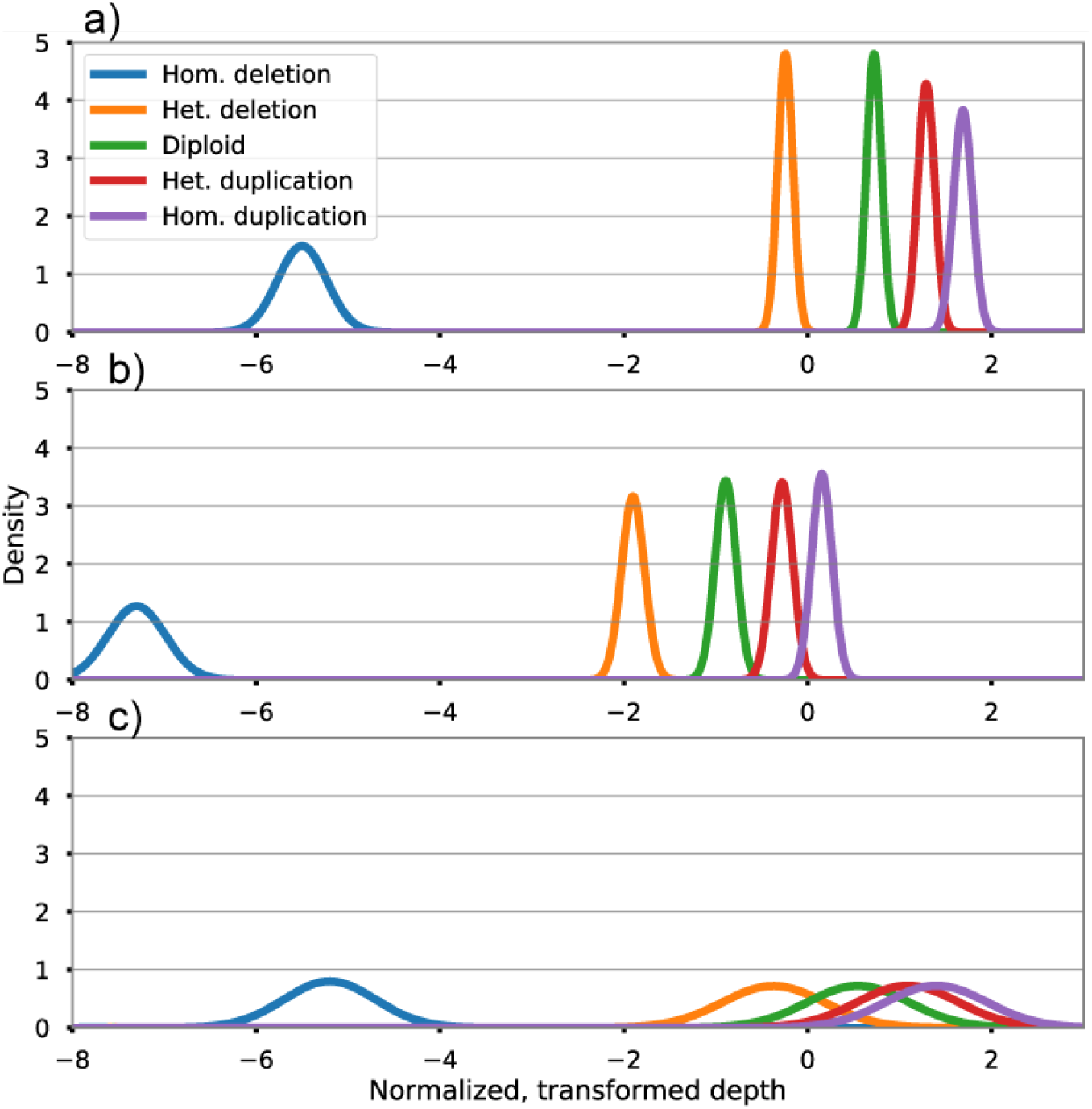
Example emission distributions illustrating between target variation. a) and b) show well behaved targets with adequate separation between emission distributions, c) demonstrates a low resolution target with substantial overlap between emission distributions.

As a proxy for overall target resolution, we compute the Kullback-Leibler divergence (Kullback & Leibler 1951) between the heterozygous deletion state and the diploid state. While the value is based only on the difference between two of the five typical states, we note that separation between the diploid and heterozygous deletion states is often closely correlated to separation between other states, and that the heterozygous deletion state is often the most relevant from a clinical standpoint. Our implementation can compute this statistic for every target in a saved model and emit the results in BED format. In practice, values less than approximately 10 appear to be associated with relatively poor CNV calling accuracy.

This procedure may be useful for understanding which targets are associated with poor resolution, and hence should be excluded from routine calling procedures. In a panel design context, such information might be used to generate formal statements regarding inclusion / exclusion of particular genes or exons.

#### 1.3.5 CNV prediction

Prediction requires a set of parameter estimates and singular vectors as computed during training, and a set of sample depths taken over the same set of targets used for model training.

We construct a homogeneous Hidden Markov Model (HMM) with initial state vector *π* = {0.01, 0.01, 0.96, 0.01, 0.01}, a transition probability matrix *M*, and set of emission distribution parameters obtained from the stored model. We construct a two-parameter variant of *M*, where one parameter describes the probability of moving ‘away’ from the diploid state and the other describes moving back ‘toward’ the diploid state. Specifically, we construct *M* as:

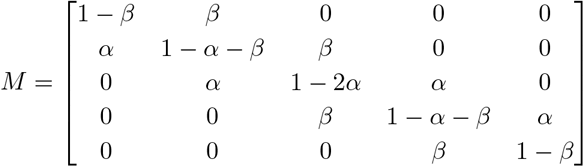

where *α* and *β* are set to 0.05 by default. Smaller values of the *α* and *β* favor fewer, larger CNVs, while larger values favor smaller, more numerous CNV calls.

Raw sample depths are transformed via *f*, and the transformed depths are treated as observations to obtain posterior state probabilities from the HMM using the Forward-Backward algorithm. Finally, we produce a list of most likely state probabilities using PMAP (pointwise maximum *a posteriori*, see Lember & Koloydenko 2013). Adjacent segments of identical most-likely state are joined into single regions, and all non-diploid regions are emitted as CNV calls.

## 2 Results

We evaluate CNV caller accuracy with a combination of simulated CNVs and those detected by array CGH and confirmed with an orthogonal technology.

### 2.1 Simulation analysis

Sensitivity and positive predictive value (PPV) were tabulated on a sample-by-sample basis given the optimal quality score cutoff for each caller and CNV size (see Table 2). A 50% reciprocal overlap (intersection-over-union / Jaccard) rule was used to determine if a simulated CNV was called correctly. Default sensitivity / specificity settings differed substantially across callers, preventing a simple comparison of raw CNV calls. To standardize caller output, we attempted to find the optimal quality score cutoff for each caller independently, then compared CNV callers using the caller-specific thresholds. For each caller candidate CNVs were called using high sensitivity settings and sensitivity / specificity curves were constructed using the caller-produced CNV call quality. The *F*_1_ statistic was calculated at 10 different evenly-spaced quality values, and the quality score associated with the maximum *F*_1_ score was chosen as the optimal quality threshold. One caller, Conifer, did not produce CNV quality scores and for this caller we did not perform quality threshold optimization.

**Table 2:** Quality thresholds used in simulation analysis

| Caller | CNV size (exons) | Quality threshold |
| --- | --- | --- |
| Cobalt | 1 | 0.95 |
|  | 3 | 0.95 |
|  | 10 | 0.96 |
| XHMM | 1 | 5 |
|  | 3 | 10 |
|  | 10 | 8 |
| ExomeDepth | 1 | -6.4 |
|  | 3 | 64 |
|  | 10 | 12 |
| Convading | 1 | 0 |
|  | 3 | 0 |
|  | 10 | 0.3 |
| Clamms | 1 | 0.85 |
|  | 3 | 0.95 |
|  | 10 | 0.93 |
| Conifer | 1-10 | NA |
| Codex | 1 | 0.52 |
|  | 3 | 30 |
|  | 10 | 118 |

**Table 3:** Number of previously detected CNVs

| Number of CNV targets | CNV type | Number of CNVs |
| --- | --- | --- |
| 1-4 | Deletion | 19 |
| 5-9 | Deletion | 14 |
| 10+ | Deletion | 5 |
| 1-4 | Duplication | 9 |
| 5-9 | Duplication | 3 |
| 10+ | Duplication | 12 |

For deletions in all size categories Cobalt demonstrated consistently high sensitivity and PPV, and achieved the highest sensitivity and PPV among all callers for deletions spanning 1-3 exons (Figure 2a & c). For deletions CNVs spanning 10 exons sensitivity was slightly higher for CODEX (Jiang et al. 2015), though at the cost of significantly lower PPV (Figure 1e). For duplications spanning 1-3 exons Cobalt demonstrated the highest mean sensitivity and PPV of the callers we examined (Figure 2b & d), although overall sensitivity for 1-exon duplications was low (61%). For 10-exon duplications, CODEX again achieved higher sensitivity (91% compared to 83%) but significantly lower PPV (70% vs 92%, Figure 2f).

**Figure 2:**
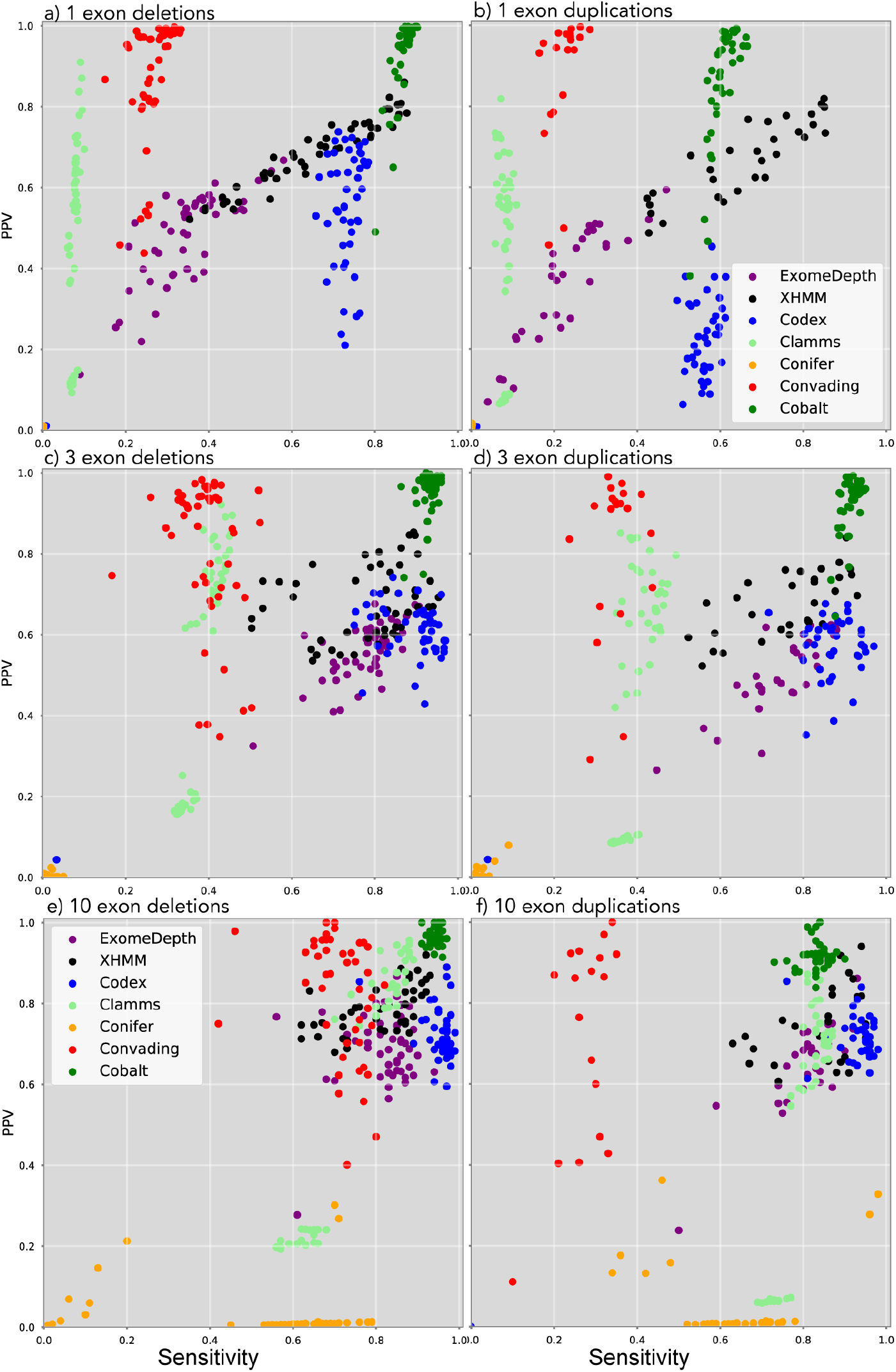
Per-sample sensitivity and positive predictive value (PPV) for simulated deletions (left column) and duplication (right column) CNVs spanning 1 (top row), 3 (middle row), and 10 (bottom row) capture targets

### 2.2 Previously detected CNVs

We examined 68 samples containing CNVs previously detected by aCGH or MLPA in addition to 160 ‘background’ samples which had not undergone any CNV detection procedure. Unlike the simulation analysis, samples were separated by sex, yielding 78 male and 82 female background samples. Sequencing and primary analysis were performed as described in the ‘NGS Data’ section. To determine if a true CNV was detected, we computed a modified Jaccard (intersection-over-union) statistic. Specifically, we computed the intersection of CNV targets in the ‘true’ CNV and the CNV targets overlapped by CNV calls, and compared this to the union of true CNV targets and the called CNV targets. If no called CNVs overlapped the true CNV a value of 0 was recorded. True CNVs with a Jaccard value of 0.5 or greater were labeled as correctly called. Caller specific quality thresholds were used as given in Table 2.

Cobalt demonstrated a sensitivity comparable to other top-performing callers, with values near 90% for small (1-4 target) deletions and near 100% for medium to large deletions (spanning 5 or more targets). For duplications, Cobalt struggled to detect small (1-4 target) CNVs and yielded a sensitivity of near 50%, but correctly detected 90-100% of medium and large duplication CNVs. CODEX achieved similar sensitivity levels for deletions and substantially higher sensitivity to small duplications, but relatively low sensitivity to large (10 or more target) duplications.

The total number of CNV calls varied substantially across callers (figure 3). While the true status of these calls is unknown, we suggest that the total number of CNV calls is positively correlated with the false discovery rate of the caller for the following reasons. First, if our sensitivity data (figures 2 and 3) are accurate, then approximately 90-100% of true CNVs are detected, thus large discrepancies in the number of CNV calls made in total are more easily explained by additional false positive calls rather than very large numbers of true, previously undetected CNVs. Second, prior analysis with aCGH has suggested that a typical number of CNVs per sample is between 0-10, and that very few samples contain hundreds (or thousands) of CNV calls.

**Figure 3:**
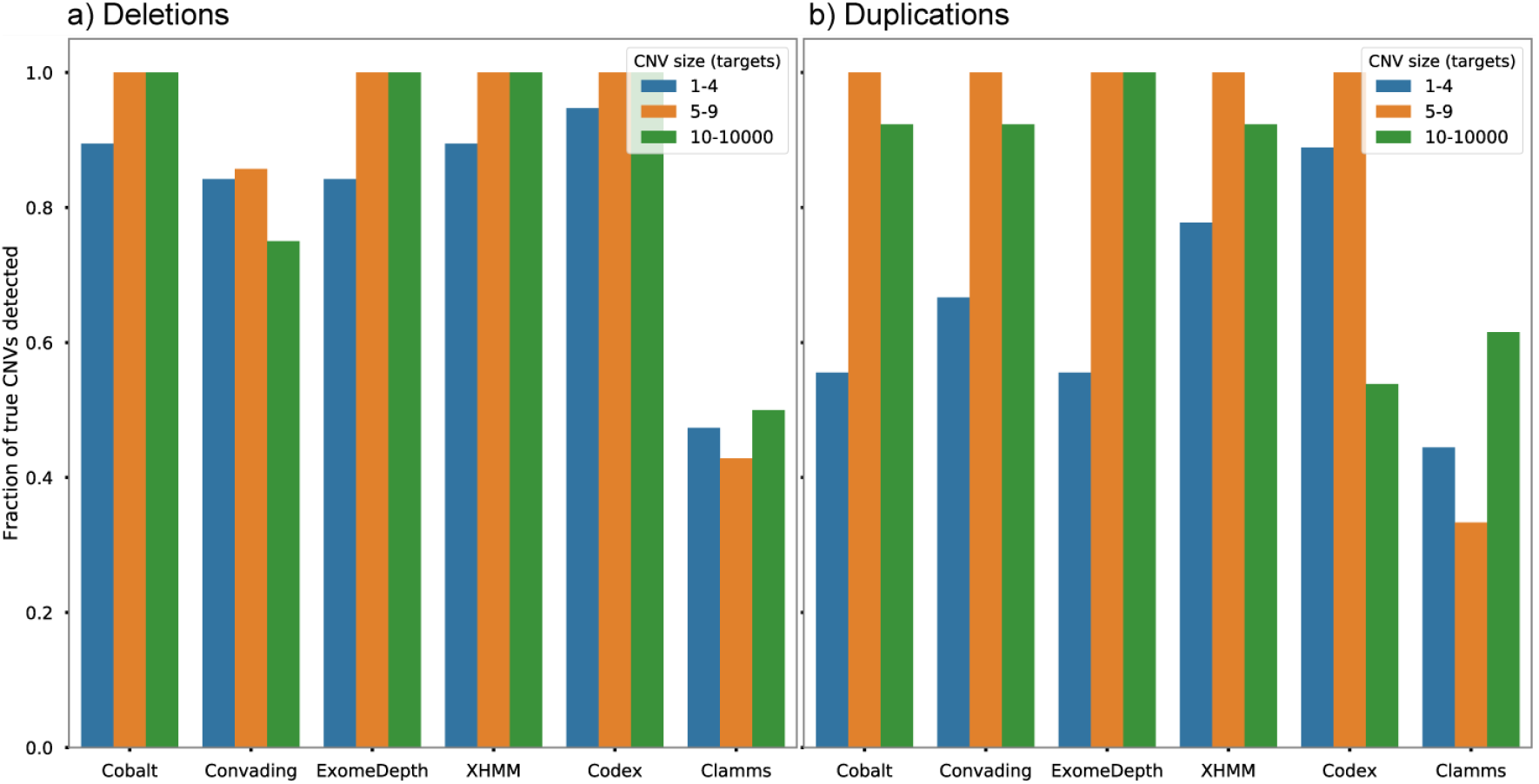
Sensitivity of Cobalt and other CNV detection tools on a) deletion and b) duplication CNVs of different sizes.

If the total number of CNV calls is correlated with false discovery rate (FDR), then Cobalt achieves the second-lowest FDR of all callers (figure 4) with a median number of 12 CNV calls per sample. The other callers that achieved similar sensitivity, CODEX, ExomeDepth, and XHMM, discovered 51, 92, and 124 calls per sample. Only Clamms yields fewer CNV calls per sample (median 2), although Clamms also demonstrated low sensitivity overall with values less than 50% in for most CNV classes.

**Figure 4:**
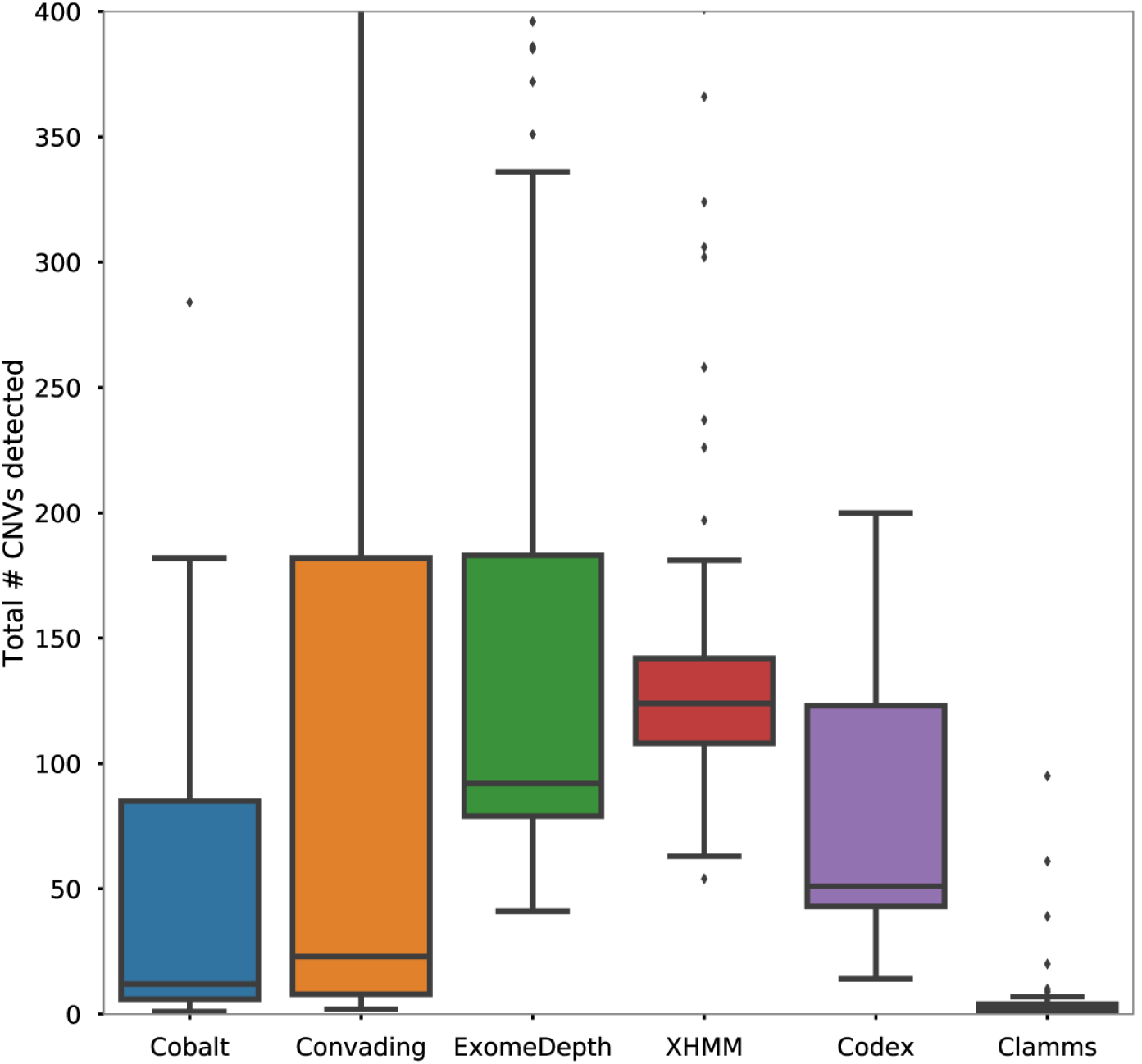
Total number of CNVs detected by Cobalt and other callers in samples with previously detected CNVs

## 3 Discussion

The Cobalt CNV caller introduces several improvements over previous techniques that improve specificity while maintaining high sensitivity. Most importantly, the HMM Cobalt uses to detect CNVs uses target-specific emission distributions. For each copy-number state at each target, these parameter estimates are produced by modifying the background sample depths to produce a pseudo-CNV, then capturing the mean and variance of the depths after log-normalization and removal of singular vectors. For comparison, the HMM of Fromer et al. (2012) assumed that the means and variances of the emission distributions were constant across all targets.

An additional refinement is use of the PMAP (pointwise maximum *a posteriori*) criterion when decoding the HMM. Most previous HMM-based CNV callers employ the Viterbi algorithm to assign copy-number states to targets. While Viterbi yields the single most likely path through states across targets, it does not necessarily maximize the number of correctly called targets, while PMAP does (Holmes & Durbin 1998). One drawback to PMAP is that it may yield *inadmissible* paths that have 0 prior or posterior probability. For instance, it may be the case that PMAP indicates a duplication target adjacent to a deletion target, even though the transition matrix specifies 0 probability for such a transition. While unsatisfying, we appeal to the approximate nature of the HMM transition matrix and note that few real-world cases justify entries of exactly 0 in the transition matrix. While it is possible to construct a PMAP decoder to yield only admissible paths (Lember & Koloykendo 2013), we leave this refinement along with exploration of more sophisticated decoding strategies to future work.

When used to predict simulated CNVs in exome data, Cobalt achieved consistently high sensitivity and PPV (Figure 2), with the exception of single exon duplications. The Codex algorithm (Jiang et al. 2015) demonstrated somewhat higher sensitivity for several categories, in particular for large duplications, but the improved detection rate comes at the cost of substantially lower PPV. Results for previously detected (‘true’) CNVs were similar, although resolution was somewhat impaired by the smaller number of true CNVs available. Cobalt yielded sensitivity indistinguishable from other top-performing callers but made far fewer CNV calls overall (figures 3, 4), strongly suggesting that Cobalt yields simultaneously high sensitivity and PPV.

PPV is particularly important in the clinical laboratory setting for several reasons. Orthogonal confirmation of putative CNV calls is often expensive and time-consuming; in fact, a primary motivation for calling CNVs from NGS data is to avoid the cost and complexity of running array-based CNV detection in addition to NGS on every sample. In the absence of routine orthogonal confirmation, labs seeking to avoid erroneous patient results labs must optimize for both sensitivity and PPV. In this setting, Cobalt may represent an appealing choice because it offers sensitivity similar to other high performing callers while making significantly fewer calls overall.

The PPVs identified in our simulation analysis may be higher than those found in real data. PPV reflects the fraction of called variants that are true positives. Our simulation analysis used relatively large numbers of CNVs (1000 single exon CNVs), for a fixed sensitivity and number of false positive calls, increasing total true CNVs will result in a higher PPV. While the true number of CNVs affecting coding exons is unknown, Conrad et al. (2010) estimated a mean of 38 per sample (his table 1), although only CNVs spanning 10 or more array probes were considered.

Generally speaking, comparisons of CNV detection tools are beset by many difficulties. First, caller performance is likely to vary substantially with features of the input data, including depth, number of targets, and the structure of variability, especially batch variability. Some callers may perform well in the face of substantial variability, while others might excel only with minimally variable data. Similarly, relative caller accuracy may shift with mean read depth. An additional factor is the amount of parameter optimization performed for each caller. In this study we have performed only minimal optimization and have relied instead on identification of optimal quality score thresholds to normalize results across callers. Nonetheless, some callers may have significantly improved relative accuracy with careful tuning of parameters.

## 4 Availability

Source code for cobalt is available via git at https://github.com/ARUP-NGS/cobalt

